# Explicitly Nonlinear Connectivity-Matrix Independent Component Analysis in Resting fMRI Data

**DOI:** 10.1101/2024.08.12.607681

**Authors:** S. M. Motlaghian, B. Baker, J. M. Ford, A. Iraji, R. Miller, A. Preda, T.G. Van Erp, V. D. Calhoun

## Abstract

Independent component analysis (ICA) is a widely used data-driven technique for investigating brain structure and function to extract intrinsic networks. However, the ability of ICA, a linear mixing model, to capture nonlinear relationships is inherently limited. While nonlinear ICA can be used to estimate nonlinear+ linear mixtures, it can be useful to study the degree to which there is nonlinearity above and beyond the widely studied linear resting networks. Here, we propose a way to divide the data into sources exhibiting linear-only or explicitly nonlinear dependencies in resting functional magnetic resonance imaging (fMRI) data. Such an approach can be very informative as it allows us to evaluate the degree to which a given network might be linear, nonlinear, or both linear and nonlinear. Here, we present an enhanced connectivity-domain ICA approach, connectivity-matrix ICA, incorporating normalized mutual information (NMI) after canceling the linear effects to measure explicitly nonlinear (EN) relationships within voxel connectivity. This integration enables the identification of brain spatial maps that exhibit pronounced explicitly nonlinear dependencies while excluding linear relationships. By eliminating linear dependencies and utilizing NMI, we discover highly structured resting networks that conventional functional connectivity methods would typically overlook. The results indicate that several maps show only linear or EN relationships, and the rest of the components display both linear and nonlinear patterns. We categorized these maps as linear-only, EN-only, and linear-EN maps. We also evaluate differences in the identified networks in a schizophrenia dataset. A significant global difference has been discovered between schizophrenia and controls in some linear-EN maps, such as the frontal lobe. Moreover, the temporal lobe and thalamus display linear group differences, while the visual and motor cortex display global differences in nonlinear relationships as their primary driver of these disparities. In sum, our findings emphasize the significance of accounting for explicitly nonlinear dependencies in functional connectivity analysis and demonstrate the effectiveness of the extended cmICA approach in revealing previously unrecognized brain dynamics.

## Introduction

Functional magnetic resonance imaging (fMRI) has emerged as a prominent approach for investigating brain function over the past few decades. Functional connectivity has been widely utilized to explore relationships among different brain regions. Existing studies of fMRI data have primarily focused on linear correlations, which have proven to be a powerful tool (Vince D. Calhoun, Liu, & Adalı, 2009; Erik Barry Erhardt et al., 2011). The ease of calculating and interpreting positive and negative correlations has made linear correlation analysis prevalent in the field, revealing intriguing patterns such as the anticorrelation of the default mode network with other networks. However, emerging research has delved deeper into the nonlinear dynamic behavior of brain networks (Motlaghian, Belger, Bustillo, Faghiri, Ford, Iraji, Lim, Mathalon, Miller, Mueller, O’Leary, Pearlson, Potkin, Preda, Erp, et al., 2022). Nonlinear relationships in the brain are observed for several reasons, such as caused by the nonlinear effects of hemodynamic responses in fMRI data, which can vary both temporally and spatially, as well as across subjects (de Zwart et al., 2009; Deneux & Faugeras, 2006; Miller et al., 2001; Obata et al., 2004). Even considering a few examples of these nonlinear effects suggests that conventional linear analysis may fail to capture relevant nonlinear relationships between distinct brain regions (Motlaghian, Belger, Bustillo, Faghiri, Ford, Iraji, Lim, Mathalon, Miller, Mueller, O’Leary, Pearlson, Potkin, Preda, van Erp, et al., 2022; Motlaghian et al., 2021).

In this work, we focus on estimating networks from explicitly nonlinear information measures. We were particularly interested in several questions. First, we want to evaluate the degree to which networks that were estimated via a linear approach were also estimable from purely nonlinear interactions. Secondly, we were interested in characterizing which networks were identified via a linear approach and were not found via a linear one or vice versa. And finally, we hypothesize that the nonlinear networks would be sensitive to group differences in schizophrenia, thus providing a new window into brain changes associated with the disorder.

Our approach combines two previously distinct lines of research: The first study is group connectivity matrix independent component analysis (cmICA) (Wu, Calhoun, Jung, & Caprihan, 2015). This study introduces a method to analyze brain connectivity, which utilizes the group ICA framework to identify resting brain networks from a large dataset. This method operates on the linear connectivity matrix to pinpoint these networks. The analysis then explores how the connectivity patterns within these resting brain networks differ across individuals. The second study used normalized mutual information (NMI), an information-theoretic, to measure linear and explicitly nonlinear dependencies between two distributions. We used this measure in our prior work (Motlaghian, Belger, Bustillo, Faghiri, Ford, Iraji, Lim, Mathalon, Miller, Mueller, O’Leary, Pearlson, Potkin, Preda, Erp, et al., 2022; Motlaghian et al., 2021) after first removing linear dependencies to assess explicitly nonlinear dependency among ICA time courses in rsfMRI data. In this work, combining the two methods, we first compute a voxel-by-voxel connectivity matrix using either linear correlation or NMI after linear canceling, then enter these matrices into a group ICA framework to estimate either linear or component maps from the resulting linear or explicitly nonlinear voxel-by-voxel connectivity matrix. By integrating these methodologies, we sought to uncover the brain regions exhibiting explicitly nonlinear relationships that are hidden when only considering linear connectivity analysis. By considering different types of spatial maps, unique EN components, unique linear components, and shared linear and EN components that are highly (spatially) similar, our objective is to comprehensively validate and investigate the various interaction patterns exhibited in the spatial domain. We then investigated the presence and extent of explicitly nonlinear (EN) relationships among brain components, then we compared them with spatial maps estimated via a linear approach, and lastly, we used our result to perform statistical comparisons of group differences between healthy controls (HC) and schizophrenia (SZ) patients.

## Participants and Preprocessing

### 2.1. Participants and Preprocessing

We used the fBIRN dataset analyzed previously used in (Damaraju et al.). The final curated dataset consisted of 160 healthy participants (mean age 36.9, 117 males; 46 females) and 151 age- and gender-matched patients with schizophrenia (mean age 37.8; 114 males, 37 females). Eyes-closed resting-state fMRI data were collected at seven sites across the United States (Keator et al., 2016). Informed consent was obtained from all subjects before scanning by the Internal Review Boards of affiliated institutions. Imaging data of one site was captured on a 3 Tesla General Electric Discovery MR750 scanner, and the rest of the six sites were collected on 3 Tesla Siemens Tim Trio scanners. Resting-state fMRI (rsfMRI) scans were acquired using a standard gradient-echo echo-planar imaging paradigm: FOV of 220 × 220 mm (64 × 64 matrices), TR = 2 s, TE = 30 ms, FA = 770, 162 volumes, 32 sequential ascending axial slices of 4 mm thickness and 1 mm skip.

Data were preprocessed by using several toolboxes such as AFNI, SPM, GIFT. Rigid body motion correction using the INRIAlign (Freire & Mangin, 2001) toolbox in SPM was applied to correct head motion. To remove the outliers, the AFNI3s 3dDespike algorithm was performed. Then fMRI data were resampled to 3 mm3 isotropic voxels. Then, data were smoothed to 6 mm full width at half maximum (FWHM) using AFNI3s BlurToFWHM algorithm, and each voxel time course was variance normalized. Subjects with large movements were excluded from the analysis to mitigate motion effects during the curation process. The preprocessing resulted in a spatial 3D by time matrix for each subject of size (53×63×52) ×157.

## Method

We introduced a new method to estimate explicitly nonlinear (EN) resting networks and compare them to the resting networks from the conventional ICA approach (Vince D Calhoun, Kiehl, & Pearlson, 2008; Vince D. Calhoun et al., 2009). The latter approach input is a time-by-voxel matrix. To assess the explicitly nonlinear relationship, which is the primary focus, we first remove the linear effect and compute the voxel-by-voxel nonlinear dependencies. Throughout this section, we explain our innovative approach, which involves applying cmICA to EN connectivity matrices. A connectivity domain (Iraji et al., 2016) approach called connectivity matrix ICA (cmICA) (Wu et al., 2015) was used to perform ICA directly on the voxel-by-voxel connectivity matrix ***C***. A modification we use is to remove the linear effects before computing ***C***, and then by NMI we estimate the explicitly linear relationships among voxels. In this case, ***C***(*i, j*) shows the explicitly nonlinear dependency between voxels *i* and *j* to identify the EN. The results derived from this methodology can be efficiently computed by without computing the full matrix ***C***, as discussed in (Wu, Caprihan, Bustillo, Mayer, & Calhoun, 2018).

### 3.1. Quantifying Explicitly Nonlinear Dependency via a Normalized Mutual Information Approach

This section explains how to assess explicitly nonlinear dependency between two distributions. Various scenarios can exist regarding their relationships for any given x and y distributions. The most commonly observed one is a linear correlation between the two. However, the relationship between these distributions can extend beyond linearity and encompass other potential cases. In this study, we employed NMI as a metric to assess relationships beyond linear correlation. We first remove the linear relationship aspect to capture the explicitly nonlinear dependencies often overlooked in conventional analyses, which are solely based on linearity. This process can be summarized into the following steps: We first estimate the linear correlation measured by a linear regression 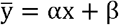, where 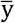 is the best linear fit predicting y given x, α is the slope, and β is the vertical intercept. Next, the linear effect is removed by calculating 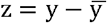. Then, the dependency between x and z is measured by NMI. Observing that this process is not symmetric when switching x and y, we defined the EN dependency of pair x and y as the average to guarantee symmetricity.

There are multiple options to standardize the mutual information, as (Kvalseth, 2017) discussed: 1) min(H(x), H(y)), 2) H(x) + H(y), and 3) max(H(x), H(y)). We employ the latter, as it has also been shown to be a (normalized) similarity metric (Horibe, 1985).

The formula we use for calculating the value of NMI is

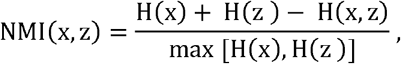

where H(x) and, H(z) are marginal entropies and H(x, z) is the joint entropy. The NMI measurement can have values between 0 and 1. If it has a value of 0, there is no dependency between x and z, and 1 indicates an absolute dependence among the two variables.

### 3.2. Linear and Explicitly Nonlinear cmICA Analysis

We embarked on utilizing the cmICA method, introduced by (Wu et al., 2015), to explore the brain’s intricate connectivity patterns. This method allowed us to extract both linear (conventional) and explicitly nonlinear spatial maps from the linear and explicitly nonlinear brain connectivity matrices. The cmICA technique facilitates the dual parcellation of brain connectivity, encompassing a set of spatially independent maps (s_k_) alongside their corresponding dual spatial maps (r_k_). These dual maps, r_k_, define the brain regions connected to the spatially independent maps, s_k_. Our primary focus in this study revolved around extracting the spatially independent maps, s_k_, from brain connectivity.

Brain functional connectivity, represented by a voxel-by-voxel matrix, provides invaluable insights into the relationships between individual voxels over time. The (i, j) entry represents the strength of the relationship between voxel i and voxel j. To comprehensively assess the dependencies between voxels, we employed two distinct metrics for calculating the connectivity matrices. Firstly, we employed Pearson correlation, a widely utilized measure for capturing linear correlations. The resulting connectivity matrix, driven by Pearson correlation, reveals the degree of linear similarity between each voxel pair. Notably, the values within this matrix ranged from -1 to 1, representing the strength and directionality of the linear relationship. In addition to Pearson correlation, we explored explicitly nonlinear dependencies using an alternative measure. This measure provides insights into the dependencies between voxels that linear correlations cannot accurately assess. The resulting connectivity matrix, reflecting explicitly nonlinear dependencies, exhibits values ranging from 0 to 1, representing the presence and strength of explicitly nonlinear relationships.

We also used a simple simulation to show possible outcomes from the EN cmICA approach. We defined voxels v_1_, v_2_, v_3,_ and v_4_ such that 1) v_1_ and v_2_ have only a linear correlation, 2) v_1_ and v_3_ have an EN and dominant linear correlation, 3) v_1_ and v_4_ have only EN relationships. After implementing linear and EN cmICA, the result showed two linear and three EN cmICA sources. We dropped v_3_ and repeated the steps. This time, the result showed two linear and two EN cmICA sources. This suggests that dominant linear correlation can overlook other voxel relations.

To identify spatial maps and explicitly nonlinear spatial maps, we first implemented the following steps for an individual dataset (Figure 1). First, the rsfMRI data, initially represented as a 3D matrix with a temporal dimension, is transformed into a vector-by-time format to facilitate subsequent analysis. This conversion involves assigning a single long dimension to each voxel for every time point. For each subject, the rsfMRI data is organized as a matrix of dimensions 53 × 63 × 52 × 157, representing the spatial dimensions (x, y, z) and time. To ensure consistent and reliable analyses, we applied a group mask by considering only voxels that contain complete brain information across all individuals.

**Figure 1.**
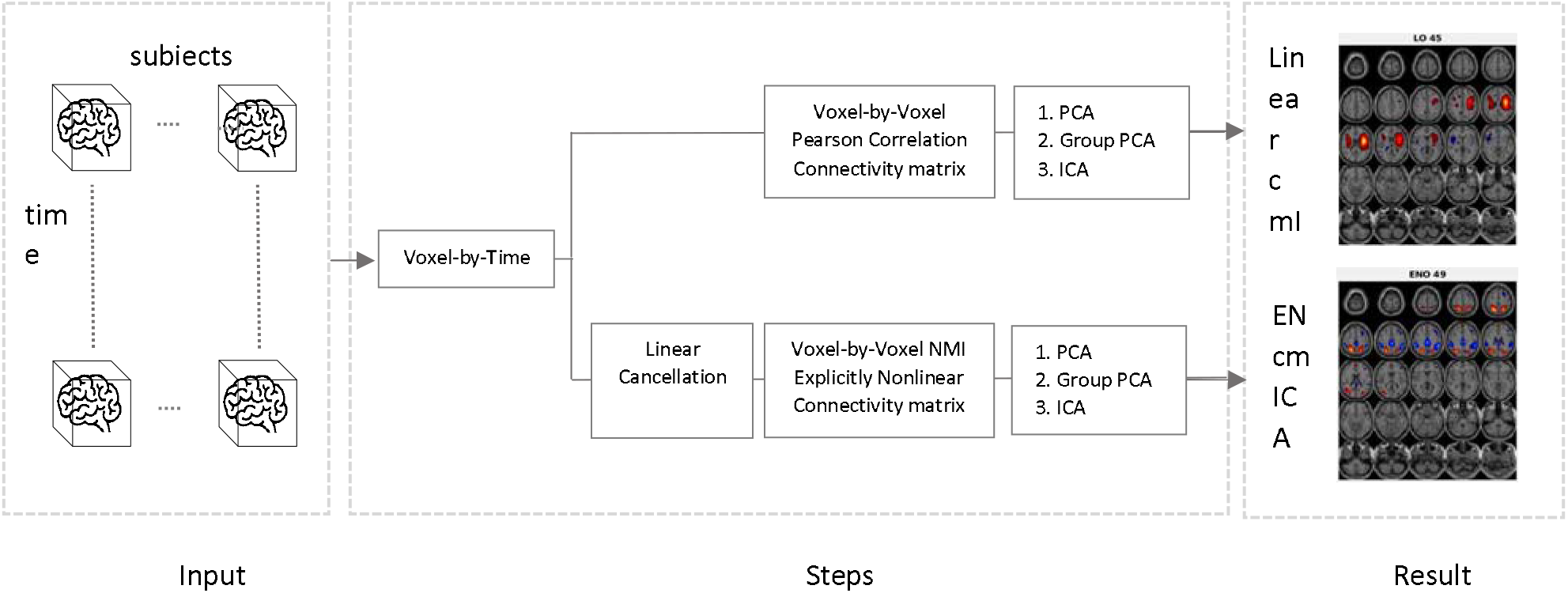
A high-level illustration of attaining linear (conventional) and explicitly nonlinear spatial maps process. The EN connectivity matrix is obtained by first canceling linear correlation for each pair of voxels and measuring the residual dependency by NMI.

Additionally, we adopted a subsampling approach to reduce computational complexity, considering every other subsample for each individual, further streamlining the subsequent analysis. This step resulted in a 5462 by 157 matrix after flattening the spatial dimensions for each individual. Next, we calculated each individual’s Pearson correlation and explicitly nonlinear (EN) connectivity matrix. These matrices are the cmICA approach’s inputs. The cmICA method thus included three main steps to identify group spatial maps: 1. PCA with 60 PCA components, 2. Group PCA with 50 PCA components and 3. ICA with 60 components. This process identified 50 linear and 50 explicitly nonlinear (EN) spatial maps. Finally, individual spatial maps were obtained using the back reconstruction method from group maps.

### 3.3. Relation between linear and Explicitly Nonlinear Spatial Maps

Linear spatial maps are commonly identified using conventional approaches. However, the inclusion of explicitly nonlinear maps in the analysis is crucial to validate and explore spatial maps that exhibit exclusively linear, exclusively explicitly nonlinear, or both types of interactions. To clarify the terminology used in this study, we define three distinct categories of spatial maps. The first category is referred to as linear-only (LO) maps, which exclusively emerge from the linear cmICA analysis. These maps capture spatial patterns characterized solely by linear relationships via Pearson correlation. The second category comprises explicitly nonlinear or EN-only (ENO) maps, which are uniquely identified through EN cmICA analysis. These maps represent spatial patterns that exhibit explicitly nonlinear interactions but are not captured by linear cmICA. Lastly, the third category encompasses the match between the linear and explicitly nonlinear (L-EN) maps. These maps represent the shared spatial patterns that can be obtained through both linear and EN cmICA analysis. They capture the presence of both linear and explicitly nonlinear interactions within the data.

Then, we manually identified components to be considered non-artifactual intrinsic networks in gray matter, not edges, ventricles, or white matter. After that, we grouped these into three categories: 1) Linear-only (LO), 2) EN-only (ENO), and 3) Linear-Explicitly Nonlinear matched (L-EN). To classify maps into the categories of LO, ENO, and L-EN, we employed a voxel-wise correlation analysis between each pair of linear and EN maps. For the L-EN category, maps with correlations exceeding 0.60 were considered as exhibiting both linear and explicitly nonlinear interactions. These maps demonstrated a strong correspondence in their spatial patterns and were indicative of shared characteristics between linear and explicitly nonlinear mappings. Maps that were uniquely identified through linear (EN) cmICA analysis are considered LO (ENO) if their absolute correlations with all EN (linear) maps are below 0.45. It is worth mentioning that we had no pair of maps with a correlation between 0.60 and 0.45.

### 3.4. Group Differences

We obtained individual cmICA maps via back reconstruction (E. B. Erhardt et al., 2011). Then, we compared the linear and explicitly nonlinear maps for two groups of HC and SZ using several methods. We also introduce a global (network level) measure to compute the significance of linear-only, EN-only, or L-EN networks. For that, the correlation between the HC maps and the SZ maps was computed using two-sample t-tests for each map. For that, we restricted the voxels to ones that were strongly present in the group map (have a z-value larger than 3). Next, the correlation values between HC and SZ are compared. We also used a voxel-wise comparison; the t-test for two samples of each voxel was used across HC and SZ’s individual cmICA maps. A full summary of all results is reported in the appendix.

## Results

4.1. This section illustrates the maps achieved from each approach. Maps are categorized into three groups: Linear-explicitly nonlinear matched (Figure *2*), linear-only (Figure 3), and explicitly nonlinear-only (Figure 4) maps.

**Figure 2.**
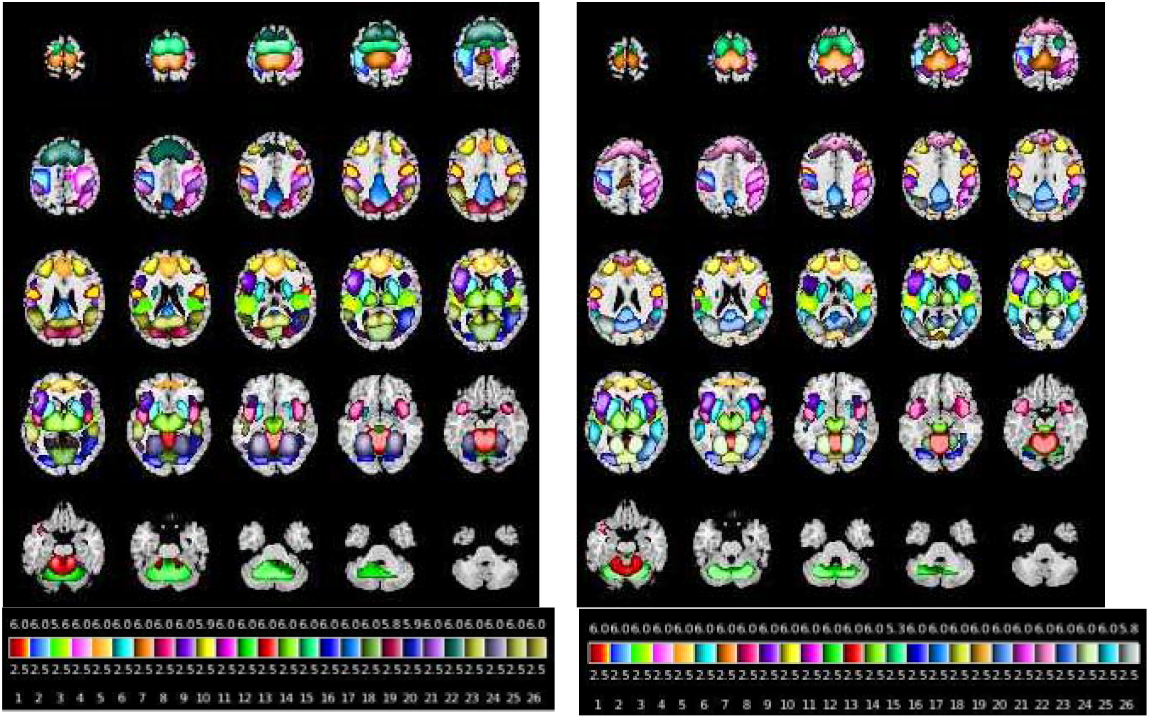
Linear (left) and EN (right) matched networks. Linear and EN maps are gained through different processes, resulting in similar networks. The left shows linear maps and the right displays the EN map. Most identified networks exhibit both linear and nonlinear properties and belong to the linear-explicitly nonlinear matched category.

**Figure 3.**
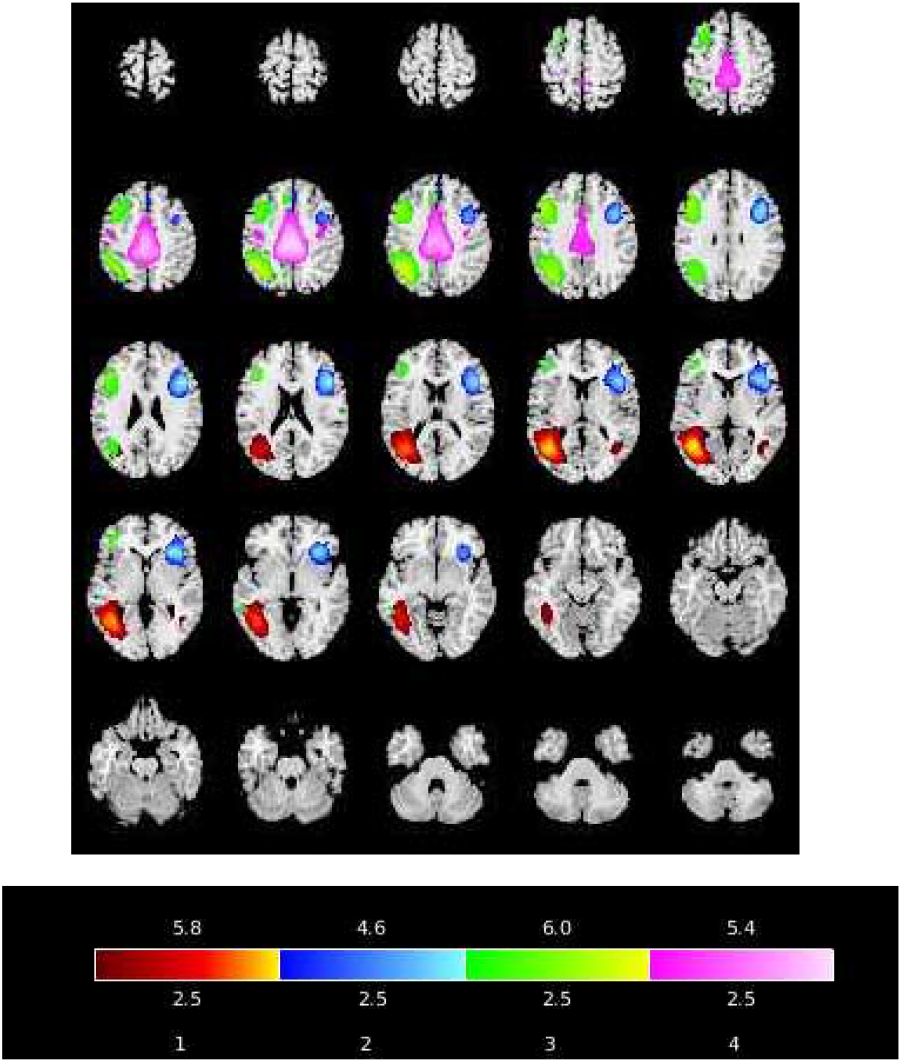
Linear-only (LO) maps. These maps are observed only in the linear cmICA results. Thus, these networks exhibit linear but not-nonlinear interactions. These networks are spread all over the brain, with some of them being unilateral networks. The left side of the brain shows more activated LO networks.

**Figure 4.**
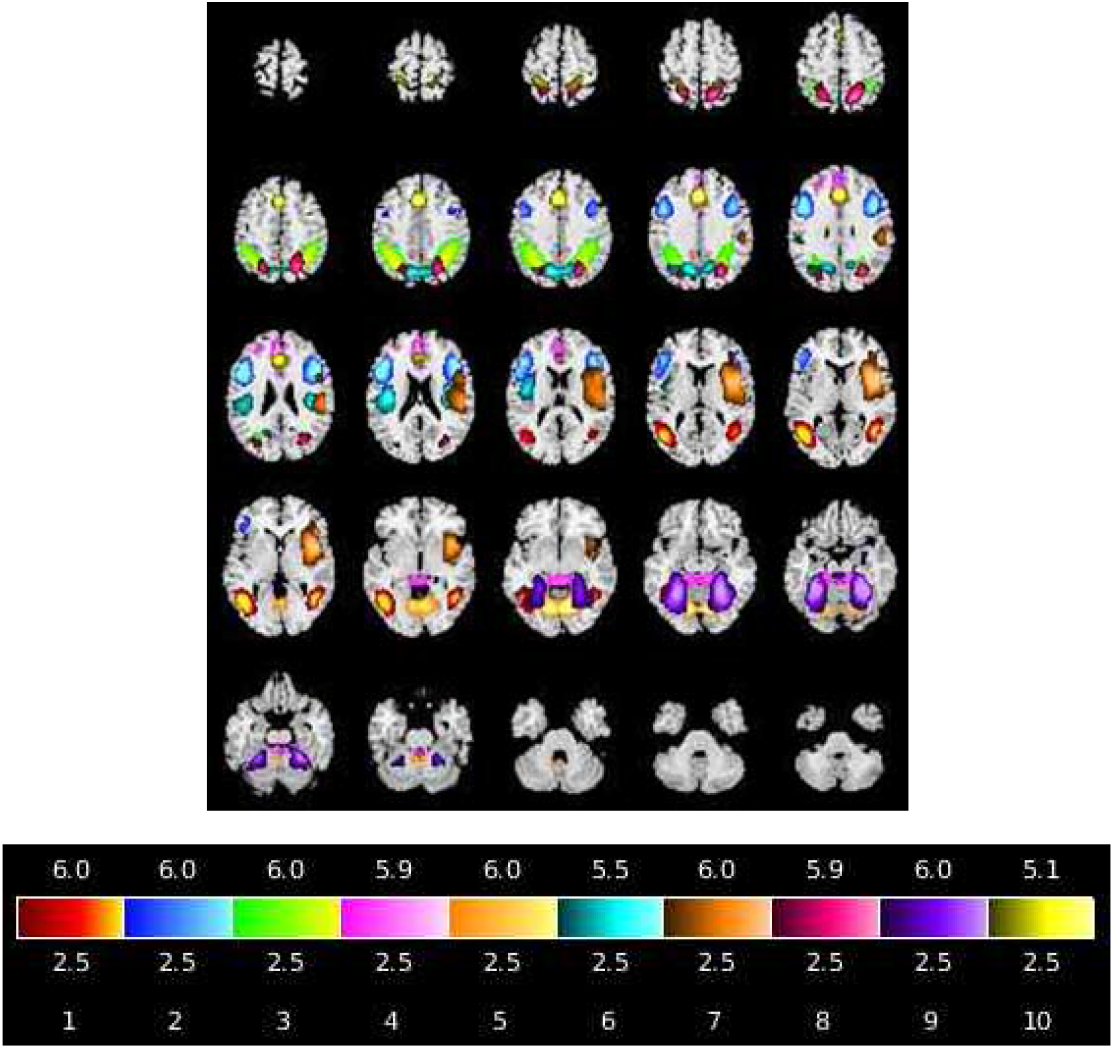
Explicitly nonlinear-only (ENO) maps. These maps are observed only in the explicitly nonlinear cmICA results. That is, they show nonlinear but not linear interactions. These maps are primarily bilateral networks. The right side of the brain shows more activated ENO networks and is not observed on the frontal or central part of the brain.

Linear and EN maps with the highest correlation are paired and displayed in Figure *2*. Our analysis identified a class of maps, “linear-explicitly nonlinear matched”, where these maps, paired one-to-one, exhibited highly similar brain regions. A strong correlation readily identified their pair for most EN and linear maps. In a few cases, more than one potential pairing emerged, and we selected the map with the stronger brain interaction for the final pairing.

Linear maps with low similarity to EN maps were assigned to a separate category, “linear-only”, displayed in Figure 3.

We created a separate category called “EN-only” maps, which showed low similarity to linear maps. These maps are then displayed in Figure *4*.

Given the complexity of the brain and fMRI data, not surprisingly, most of the networks display both linear and EN activity. We also find several networks that demonstrate linear-only or EN-only behavior. We make the following observations:

1. All EN-only maps except one are symmetric, but none of the linear-only maps are symmetrical concerning the right and left. Thus, we see a split of maps into lateralized parts in linear-only vs bilateral in EN-only.
2. EN-only maps cover most of the brain, from subcortical to cortical, while linear-only maps are primarily in subcortical regions or the left hemisphere.
3. There were more EN maps than linear maps overall, suggesting the majority of the brain exhibits nonlinear effects.

### 4.2. Group Differences

To further investigate group differences between healthy controls (HC) and schizophrenia (SZ) patients, global measures were computed for both linear and EN maps. Notably, all the linear and EN maps that show a significant difference belong to the linear-EN group, i.e., none of the EN-only or linear-only showed significant differences (Figure 5 and Figure 6). While the frontal lobe and right-lateralized motor networks demonstrated significant group differences in both linear and EN domains (Figure 6), other regions displayed significant differences in either the linear (Figure 5 left) or EN (Figure 5 right) domain exclusively. The temporal lobe and thalamus exhibit primarily linear group differences, whereas the visual cortex and left motor cortex reveal group differences exclusively linked to nonlinear patterns. This suggests that schizophrenia is associated with either linear or nonlinear changes in specific brain regions.

**Figure 5.**
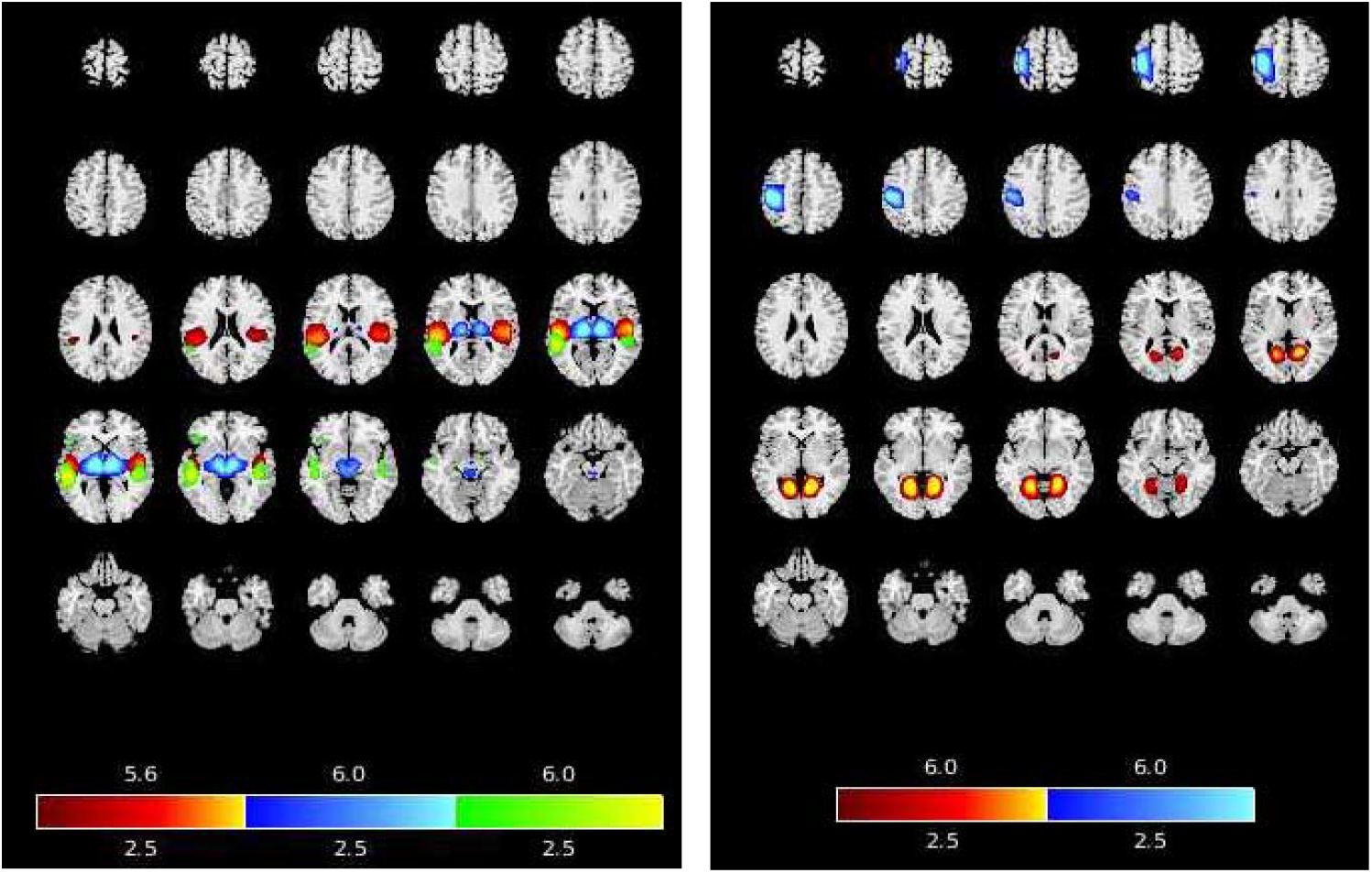
Linear networks showing group differences (left) and nonlinear networks showing group differences (right): The temporal lobe and thalamus are some of the critical areas that are expected to show differences in schizophrenia. These regions show significant linear effects and do not show significant nonlinear effects. Thus, the group differences in these networks are dominated by linear effects. In contrast, the visual cortex and left motor cortex show group differences only for the EN networks, suggesting that nonlinear relationships drive the differences.

**Figure 6.**
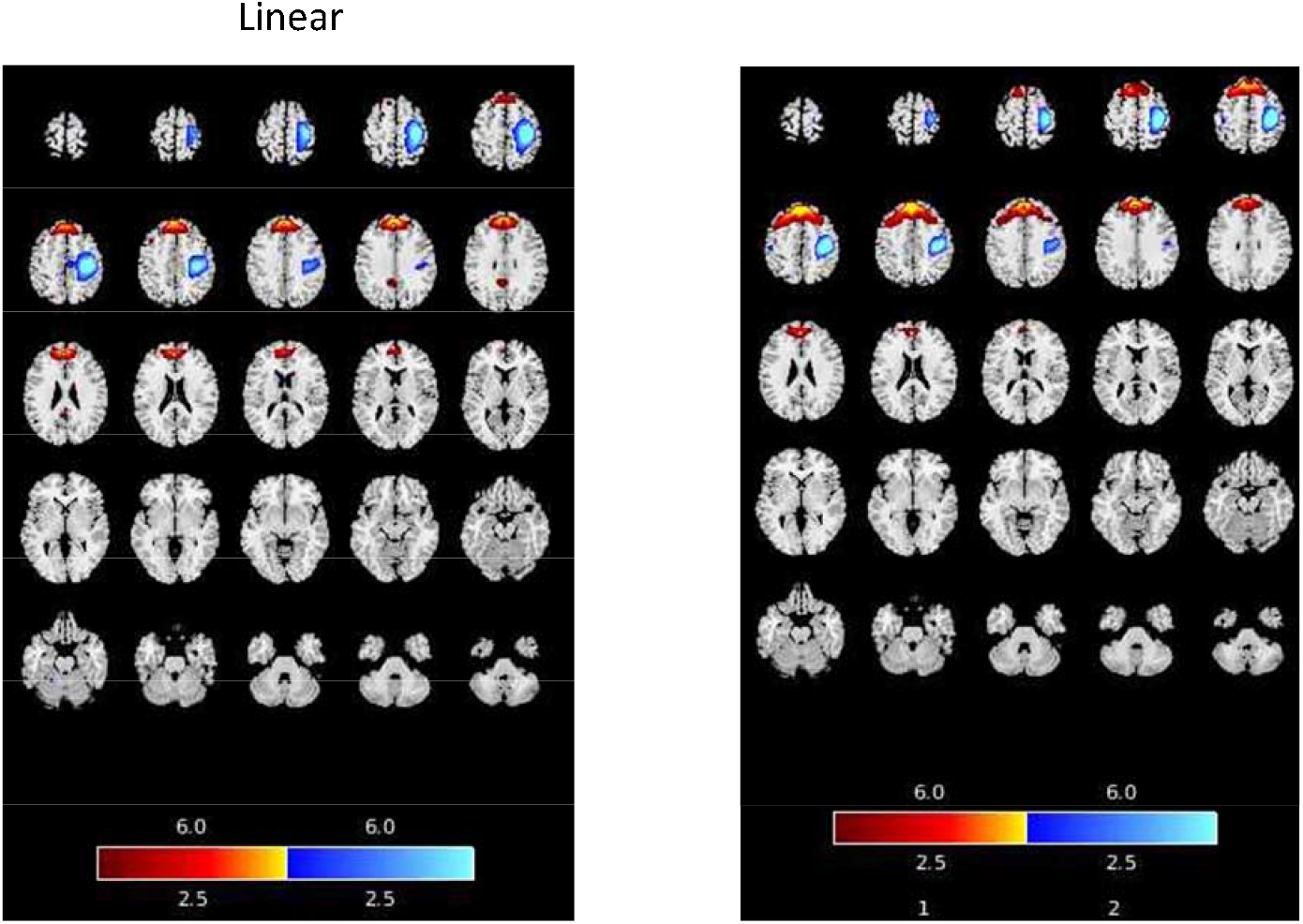
Linear and nonlinear group differences: Frontal lobe and right lateralized motor show group differences in both linear and explicitly nonlinear networks.

## Conclusion and Discussion

In this paper, we implemented a new method, EN cmICA, to identify brain networks that may otherwise be ignored or distorted by the traditional linear approach. This method is a combination of the group cmICA and NMI. Canceling the linear effect before estimating the networks by cmICA allows us to discover explicitly nonlinear networks, above and beyond the dominant linear relationships, that may show a more complex interaction with the rest of the brain. The innovation of implementing cmICA rather than ICA on the time-series data was a critical step as it allowed us to extract networks from connectivity matrices derived from mutual information after removing the linear effects.

In linear cmICA, given a voxel, a corresponding column represents its correlation with other voxels in the correlation connectivity matrix. Linear cmICA decomposes the connectivity matrix into maximally independent components, each representing a spatial map or brain network. In EN cmICA, a similar concept applies, except we use NMI instead of the correlation connectivity matrix after removing the linear relationships from the time series. The values of this matrix thus represent explicitly nonlinear relationships. Each component network extracted is thus derived from the explicitly nonlinear information. By comparing the EN networks with the linear networks, we can identify cases where similar networks are identified using only linear information, nonlinear information, or both.

Most driven networks with EN cmICA analysis were similar to the linear networks; however, a few were unique. The simulation revealed that one of the cases in which a unique EN network can exist is in an event where a dominant linear correlation coats an explicitly nonlinear relationship. Another case to have a unique network is lateraled maps that are separated in linear approach but are considered the same network in EN cmICA. Implementing EN cmICA combined with linear cmICA facilitates the discovery of lateralized brain regions. This highlights that linear and cmICA are complementary as each evaluates the brain activity through a different lens.

The global measure showed significant differences can exist either in one or both approaches for maps that can be obtained from both methods, i.e., EN cmICA maps showed group differences while the same pair in linear did not and vice versa. This also emphasizes that linear and EN approaches are needed to expand our insight into the brain’s workings. Each linear or EN cmICA approach reveals a map set that mostly overlaps; however, some maps are uniquely gained from only one method, providing a different insight into brain networks.

One of the limitations of the EN cmICA method is that the calculation of explicitly nonlinear relationships after deletion of linear correlation is costly as the process is done voxel by voxel. One may start from particular linear networks and study their EN cmICA segmentations to drive the underlined explicitly nonlinear relationship.

In sum, a new method, EN cmICA, is introduced to identify brain networks that traditional linear approaches might miss. EN cmICA combines group cmICA with techniques to remove linear trends, allowing it to capture explicitly nonlinear relationships within the brain. By comparing networks identified by EN cmICA and linear cmICA, researchers can find networks driven solely by linear or nonlinear information or by both. This highlights that both methods provide complementary insights. While computationally expensive, EN cmICA offers a valuable tool for expanding our understanding of brain function.

## Appendix A

### Supplementary Data

Supplementary data to this article can be found online at http://dx.doi.org/10.1016/j.nicl.2014.07.003

### Funding sources

This study was funded in part by NIH grants R01MH118695, R01MH123610, and R01EB006841 and NSF grants 2112455, 2316421.

